# CRISPResso: sequencing analysis toolbox for CRISPR genome editing

**DOI:** 10.1101/031203

**Authors:** Luca Pinello, Matthew C. Canver, Megan D. Hoban, Stuart H. Orkin, Donald B. Kohn, Daniel E. Bauer, Guo-Cheng Yuan

## To the Editor

Recent progress in genome editing technologies, in particular the CRISPR-Cas9 nuclease system, has provided new opportunities to investigate the biological functions of genomic sequences by targeted mutagenesis [1-4]. Briefly, Cas9 may be directed by a chimeric single guide RNA (sgRNA) to a target genomic sequence upstream of a protospacer adjacent motif (PAM) for cleavage. Double strand breaks (DSBs) resulting from site-specific Cas9 cleavage can be resolved by endogenous DNA repair pathways such as non-homologous end joining (NHEJ) or homology-directed repair (HDR). These repair mechanisms result in a spectrum of diverse outcomes including insertions, deletions, nucleotide substitutions, and, in the case of HDR, recombination of extrachromosomal donor sequences [1-3, 5, 6]. Deep sequencing of amplified genomic regions or whole genome sequencing (WGS) allows for quantitative and sensitive detection of targeted mutations. However, to date no standard analytic tool has been developed to systematically enumerate and visualize these events, resulting in inconsistencies between different experiments and across laboratories. Challenging issues for the interpretation of CRISPR-Cas9 edited sequences include amplification or sequencing errors, experimental variation in sequence quality, ambiguous alignment of variable length indels, deconvoluting mixed HDR/NHEJ outcomes, analyzing large datasets from WGS experiments and analyzing pooled experiments where many different target sites are present in a single sequencing library. To solve these issues with an aim to standardize data analysis, we developed CRISPResso as a robust and easy-to-use computational pipeline (**Supplementary Note 1** and **Supplementary Fig. 1**). CRISPResso enables accurate quantification and visualization of CRISPR-Cas9 outcomes, as well as comprehensive evaluation of effects on coding sequences, noncoding elements and selected off-target sites.

CRISPResso is a suite of computational tools that provides an integrated, user-friendly interface that can be operated by biologists and bioinformaticians alike (**Supplementary Figs. 1 and 2**). Compared to existing tools [7], CRISPResso offers several novel features, including: batch sample analysis via command line interface, integration with other pipelines, tunable parameters of sequence quality and alignment fidelity, discrete measurement of insertions, deletions, and nucleotide substitutions (ignored by other methods), tunable windows around the cleavage site to minimize false positive classification, quantification of frameshift versus in-frame coding mutations, and distinction between NHEJ, HDR, and mixed mutation events. CRISPResso automates the following steps: 1. filtering low quality reads, 2. trimming adapters, 3. aligning the reads to a reference amplicon, 4. quantifying the proportion of HDR and NHEJ outcomes, and 5. determining the proportion of frameshift and in-frame mutations as well as detecting potential splice site mutations. A graphical report is generated to visualize mutagenesis profiles (**Fig. 1**, **Supplementary Figs. 3-5**), and plain text output files are also produced for further integrative analyses (**Supplementary Note 2**). This pipeline can be used for assessment of on-target editing efficacy as well as of off-target editing at selected loci [8, 9].

**Figure 1:**
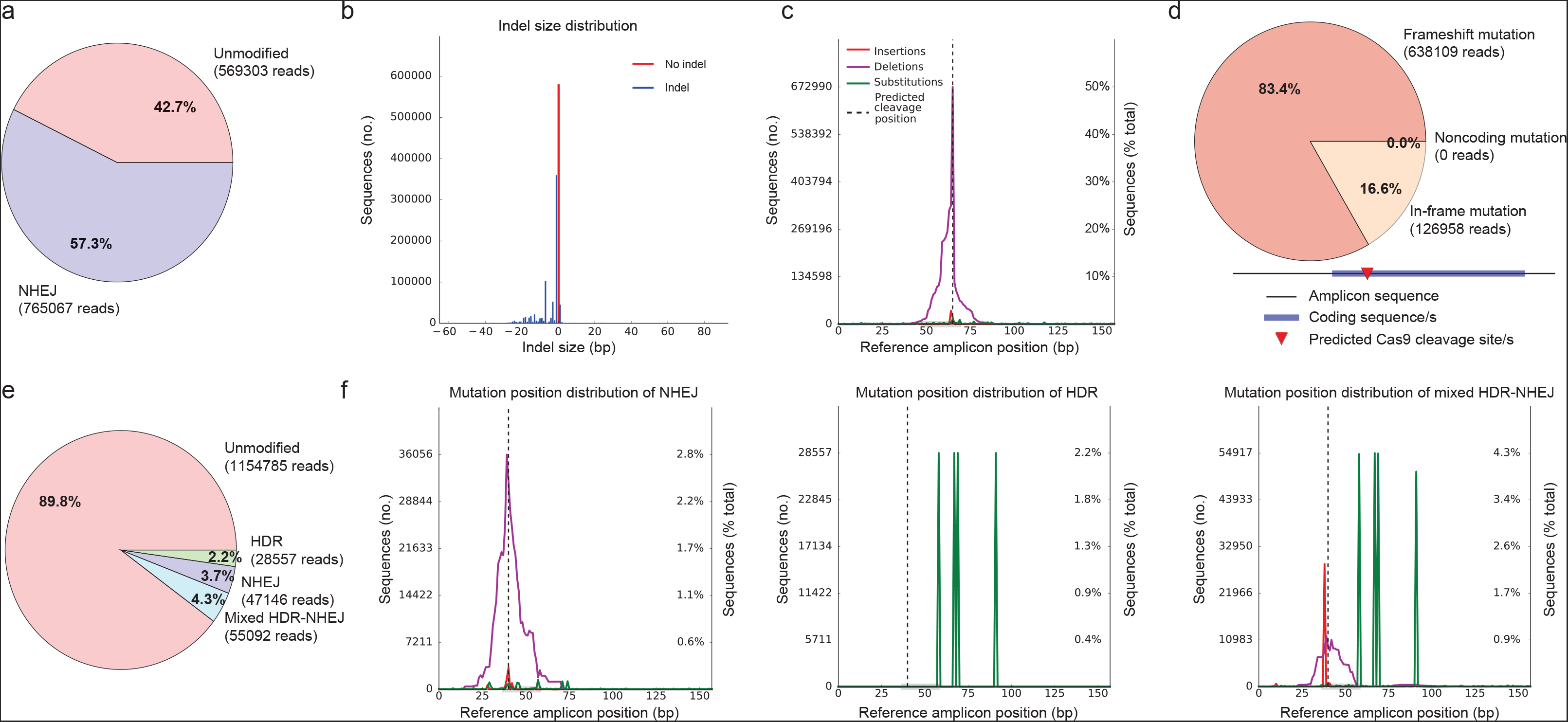
Quantification and visualization of NHEJ and HDR mutagenesis profiles. **a-d**, An example of NHEJ-mediated disruption of a coding sequence by CRISPR-Cas9 (experiment 1). **a**, Quantification of editing frequency as determined by the percentage and number of sequence reads showing unmodified and modified alleles. When no donor sequence is provided, CRISPResso classifies any mutation overlapping a window around the expected cleavage site/s as an NHEJ event. **b**, Frequency distribution of alleles with indels (shown in blue) and without indels (in red). Length-conserving substitutions are not classified as indels in this plot. In this example, the indels are dominated by small deletions, consistent with the anticipated CRISPR-Cas9 effect. **c**, NHEJ reads with insertions (red), deletions (purple), and substitutions (green) mapped to reference amplicon. For insertions, the positions immediately adjacent to the insertion are indicated. In this example, the mutations cluster around the predicted cleavage position, consistent with the anticipated CRISPR-Cas9 effect. A low level of substitutions apparent throughout the amplicon suggests low-level technical error, although these errors do not contribute to the quantification of the NHEJ. **d**, Frameshift analysis of coding sequence reads affected by modifications. Frameshift and in-frame mutations include any mutations that partially or fully overlap coding sequences as input by the user, with any non-overlapping mutations classified as noncoding (see also Supplementary Fig. 11). **e-f**, An example of HDR-mediated recombination of an extrachromosomal donor sequence resulting in four substitutions relative to the reference amplicon (experiment 2)**. e**, When an expected HDR amplicon is provided, CRISPResso classifies sequence reads as HDR if they preferentially align to the expected HDR amplicon sequence and NHEJ (or unmodified) if they preferentially align to the reference amplicon. An alignment threshold may be provided to distinguish HDR alleles from those showing evidence of mixed HDR-NHEJ repair. **f**, Mapping of mutation position to reference amplicon of reads classified as NHEJ (left), HDR (center), and mixed HDR-NHEJ (right). In this example with the alignment threshold set to 100% sequence identity, the HDR alleles show only the four expected substitutions (see Supplementary Fig. 4) while the mixed HDR-NHEJ alleles show additional indels at the predicted cleavage position, consistent with sequential cleavages initially repaired by HDR and subsequently by NHEJ.

CRISPResso qualitatively and quantitatively evaluates the outcomes of genome editing experiments in which target loci are subject to deep sequencing. We initially assessed the performance and limitations of CRISPResso, performing simulations with various genome editing outcomes, with and without sequencing errors included (**Supplementary Note 3**, **Supplementary Figs. 6-9**). We found that CRISPResso, even in the presence of sequencing errors, robustly and accurately recovers editing events with a negligible false positive rate (<=0.1%). Then we applied CRISPResso to actual paired end deep sequencing data from cells expressing Cas9 and sgRNA-1 targeted to a coding sequence with the intent to create gene knockout by frameshift mutations (experiment 1) or cells expressing Cas9, an extrachromosomal homologous donor template, and sgRNA-2 (experiment 2) or sgRNA-1 (experiment 3) with the intent of targeted introduction of four nucleotide substitutions (**Supplementary Note 4** and **Supplementary Figs. 3-5, 10**). For experiment 1, CRISPResso provides a quantification of the proportion of NHEJ occurrences, mutated allele size distribution and precise mutation localization with respect to the reference amplicon (**Fig. 1a-c**). When coding sequences are provided as an optional input, the software quantifies frameshift and in-frame mutations as well as predicts splice site mutations (**Fig. 1d, Supplementary Fig. 11**). When an expected HDR amplicon sequence is provided (experiments 2 and 3), CRISPResso is able to deconvolve and characterize unmodified, NHEJ-modified, and HDR-modified alleles as distinct outcomes (**Fig. 1e, f, Supplementary Note 1**, **and Supplementary Figs. 3-5**). In addition, it identifies mixed alleles that may result from sequential cleavages initially resulting in HDR and later NHEJ repair (**Fig. 1f**). In a case when the donor sequence disrupts the guide RNA seed sequence or PAM, the relative fraction of mixed events appears substantially reduced, consistent with the effect of these HDR alleles on resisting subsequent cleavage (**Supplementary Figs. 12, 13**). By specifying the sequence identity required to classify an event as HDR, the user can control the specificity of HDR and sensitivity of mixed HDR-NHEJ allele detection (**Supplementary Figs. 12, 13**). CRISPResso can be run either as a stand-alone command line utility (**http://github.com/lucapinello/CRISPResso**) or web application (**www.crispresso.rocks, Supplementary Note 2**).

In addition, the CRISPResso suite accommodates single or pooled amplicon deep sequencing and WGS datasets and allows the direct comparison of individual experiments. In fact, four additional utilities are provided (Supplementary Fig. 1): (1) CRISPRessoPooled is a tool for the analysis of pooled amplicon experiments that first preprocesses the input data to highlight and remove PCR amplification or trimming artifacts. This tool is recommended to be run in a mixed mode with alignment to both a reference genome and a list of amplicons to help resolve alignment artifacts or contamination (although it may also be run in individual modes). The outputs of CRISPRessoPooled include individual CRISPResso reports with detailed mapping statistics for each region as well as a summary table for all target regions (**Supplementary Note 5**, **Supplementary Figs. 14-15** and **Supplementary Tables 1-6**). (2) CRISPRessoWGS is a tool for the analysis of WGS data that provides detailed CRISPResso reports for any set of sites throughout the genome (for example, potential off-target sites) and separate .bam files (for discrete visualization in a genome browser) (**Supplementary Note 6** and **Supplementary Fig. 16**). (3) CRISPRessoCompare is a tool for the comparison of two CRISPResso analyses, useful for example to compare treated and untreated samples or to compare different experimental conditions (**Supplementary Note 7** and **Supplementary Figure 17**). (4) CRISPRessoPooledWGSCompare is a tool to compare experiments involving several regions analyzed by either CRISPRessoPooled or CRISPRessoWGS (**Supplementary Note 8** and **Supplementary Table 7**). In summary, the CRISPResso suite offers flexible and powerful tools to evaluate and quantitate genome editing outcomes from sequencing experiments, and for standardizing and streamlining analyses that currently require development of custom in-house algorithms.

